# Identification and characterization of a *Streptococcus* strain isolated from the esophagus of an adult male heterozygous for the CF allele

**DOI:** 10.64898/2026.11.15.765035

**Authors:** E. Axel Abrahamson, Sarvesh V. Surve, George A. O’Toole

**Author notes:** Address correspondence to: George A. O’Toole, Dept. of Microbiology & Immunology, Rm 202 Remsen Building 66 College Street, Geisel School of Medicine at Dartmouth Hanover, NH 03755.

## Abstract

The genus *Streptococcus* consists of diverse bacteria that are integral components of the human microbiome, acting as both important commensals and opportunistic pathogens. In this study, one strain of an α-hemolytic species of the genus *Streptococcus*, GP0011^T^, isolated from an esophageal aspirate collected during an endoscopy, was characterised phenotypically to establish its taxonomic status. The strain was isolated from an adult male heterozygous for a *cystic fibrosis transmembrane conductance regulator* (*CFTR*) mutation. The phylogenetic analysis indicates that the strain is a novel species within the mitis group, with *Streptococcus oralis* ATCC 35037^T^ being the closest relative with 94.84% average nucleotide identity (ANI) and 58.3% digital DNA-DNA hybridization (dDDH). Phenotypic characterization and API20 Strep strips distinguished GP0011^T^ from *S. oralis* by the absence of alkaline phosphatase activity, trehalose fermentation, and starch fermentation, and by superior growth on D-mannitol and D-sorbitol, consistent with genomic data. The strain is Gram-positive, catalase-negative, and the cells are ovoid with a mean cell diameter of 1.3 µm. Growth occurred at 25-43 °C, with an optimum at 30°C, with an optimum at pH 8.0. Antibiotic resistance genes *msrD* and *mefA,* conferring resistance to azithromycin and erythromycin, as well as pathways for mannitol and sorbitol metabolism, were found and phenotypically confirmed in the GP0011^T^ genome but were absent from *S. oralis*. These findings indicate that GP0011^T^ represents a novel species within the genus *Streptococcus* with the proposed name of *Streptococcus ruoffae* sp. nov. The type strain is GP0011^T^ (ATCC TSD-627 and NCTC 15312).

## INTRODUCTION

The genus *Streptococcus* is a part of the family *Streptococcaceae*, order *Lactobacillales*, and class *Bacilli*. They are Gram-positive, catalase-negative, non-motile, facultatively aerobic firmicutes that form linear chains [1]. *Streptococcus* species inhabit the human body as opportunistic pathogens and commensal organisms, colonizing mucosal surfaces of the upper respiratory tract, gastrointestinal system, and genitourinary system [2]. Phylogenomic analyses have established two primary clades, mitis-suis and pyogenes-equinus-mutans, further subdivided into fourteen distinct subclades. Under the classification system proposed in 2018 by Patel and Gupta, the isolate described in this study falls into the mitis-suis clade and the pneumoniae subclade [3]. As of February 2026, the genus contains 141 taxa with a validly published and correct name [4].

The esophagus harbors a wide variety of microorganisms. In healthy individuals, *Streptococcus* is one of the predominant genera and *Streptococcus mitis* constitutes the most abundant species of the esophageal microbiome [5]. In patients with esophagitis and related local inflammatory and metaplastic states, however, the relative abundance of *Streptococcus* spp. decreases and is replaced by a shift toward Gram-negative anaerobes and microaerophiles [6, 7].

Cystic fibrosis (CF) is an autosomal recessive disorder caused by a mutation in the *CFTR* gene, which encodes a chloride channel important for mucosal hydration across epithelial surfaces [8]. The microbial gut dysbiosis in CF is well-established, with alterations to the diversity and composition of communities, ultimately contributing to an inflammatory response [9–11]. Heterozygous carriers have historically been considered clinically unaffected, but subtle alterations in mucosal ion transport and mucus composition have been documented as increasing risk for respiratory infections as well as many of the sequelae facing homozygotes [12–14].

Here, we report the characterization of a novel α-hemolytic *Streptococcus* isolated from the esophageal luminal fluid of a *CFTR* heterozygous patient. Based on phenotypic and genomic data, we propose the name *Streptococcus ruoffae* sp. nov. for this taxon.

## RESULTS

### Phylogenomic Analyses Establishes GP0011^T^ as a New Species

GP0011^T^ was identified through isolate collection of a clinical endoscopy sample in which it could not be assigned to any species (15). This flagged the isolate as a potential novel species. The draft genome of GP0011^T^ had 100% completeness with 0% contamination for a total genome size of 2.1 Mb, composed of 20 contigs with a contig N_50_ of 274.2 kb and L_50_ of 3 (Table 1). A total of 1984 protein-coding sequences were predicted along with 3 rRNA and 42 tRNA genes. A whole-genome phylogeny generated using TYGS with the Genome BLAST Distance Phylogeny (GBDP) method identified the most closely related type strains to GP0011^T^ (Fig. S1a, supplementary data) [4, 16]. The most similar genomes to GP0011^T^ were *Streptococcus oralis* ATCC 35037^T^ (58.3%), followed by *S. jiangnanensis* (56.6%) and *S. halitosis* (50.8%) (Fig. S1b, supplementary data), all of which fall below the species threshold of 70% and therefore suggest the presence of a distinct species from those previously classified [17].

**Table 1.**
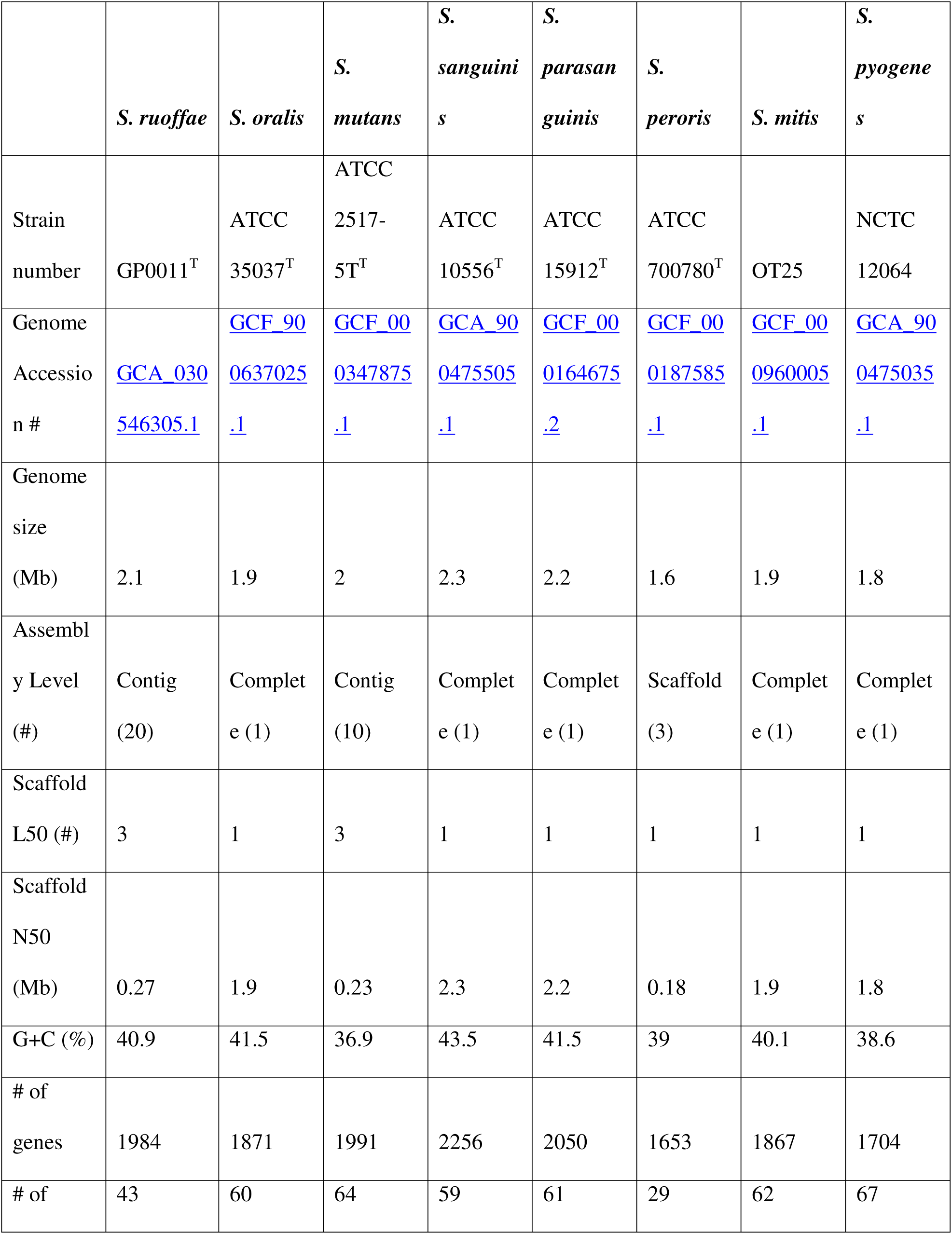
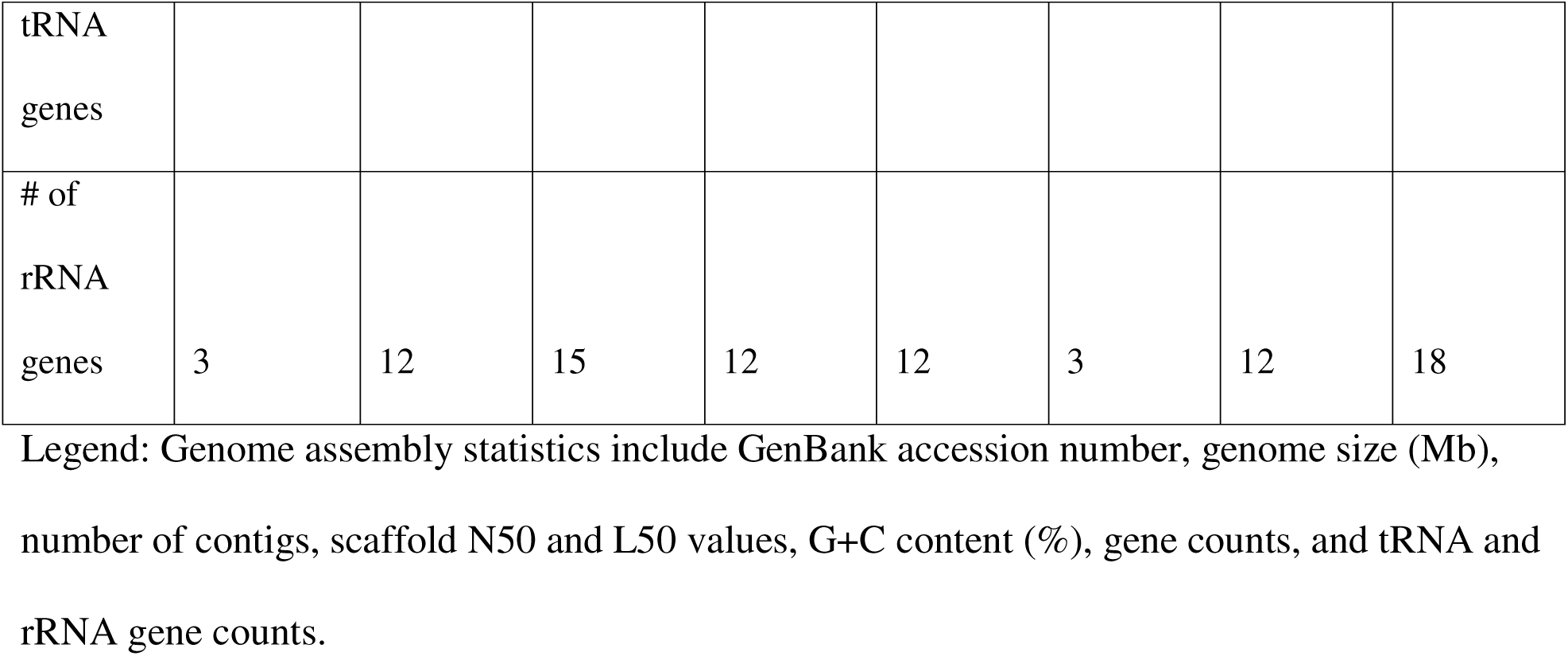
Genomic features of *Streptococcus ruoffae* GP0011.

From this analysis, and apart from *S. oralis*, the next highest shared identity was seen for *S. mitis* with ANI=86.2% and dDDH=32%, values falling far below the respective species delineation thresholds. Sequence comparison indicated that GP0011^T^ had the greatest 16S rRNA gene sequence similarity to *S. mitis* (99.42%), followed by *Streptococcus oralis* ATCC 35037^T^ (99.23%) and *S. sanguinis* (97.98%) (Fig. S2a, supplementary data) [18]. The 16S rRNA gene classification of GP0011^T^ was within the mitis group.

The GP0011^T^ *gyrB* (1946 bp) and *rpoB* (3611 bp) gene sequences were compared with related species, and their closest relatives, *S. oralis*, *S. mitis*, and *S. peroris*, showed 97.04%, 87.33%, and 83.81% respective sequence identity to *gyrB*, and 97.26%, 93.77%, and 90.17% respective sequence identity to *rpoB*, consistent with species-level divergence (Fig. S2b-c, supplementary data) [18]. A complementary maximum likelihood phylogenomic tree reconstructed for the eight *Streptococcus* species using 673 single-copy core genes aligned with MAFFT using the LG+F model in RAxML placed GP0011^T^ within the mitis group but as a distinct, well-supported lineage separate from the mitis group type strains (Fig. 1b) [19, 20].

**Fig. 1.**
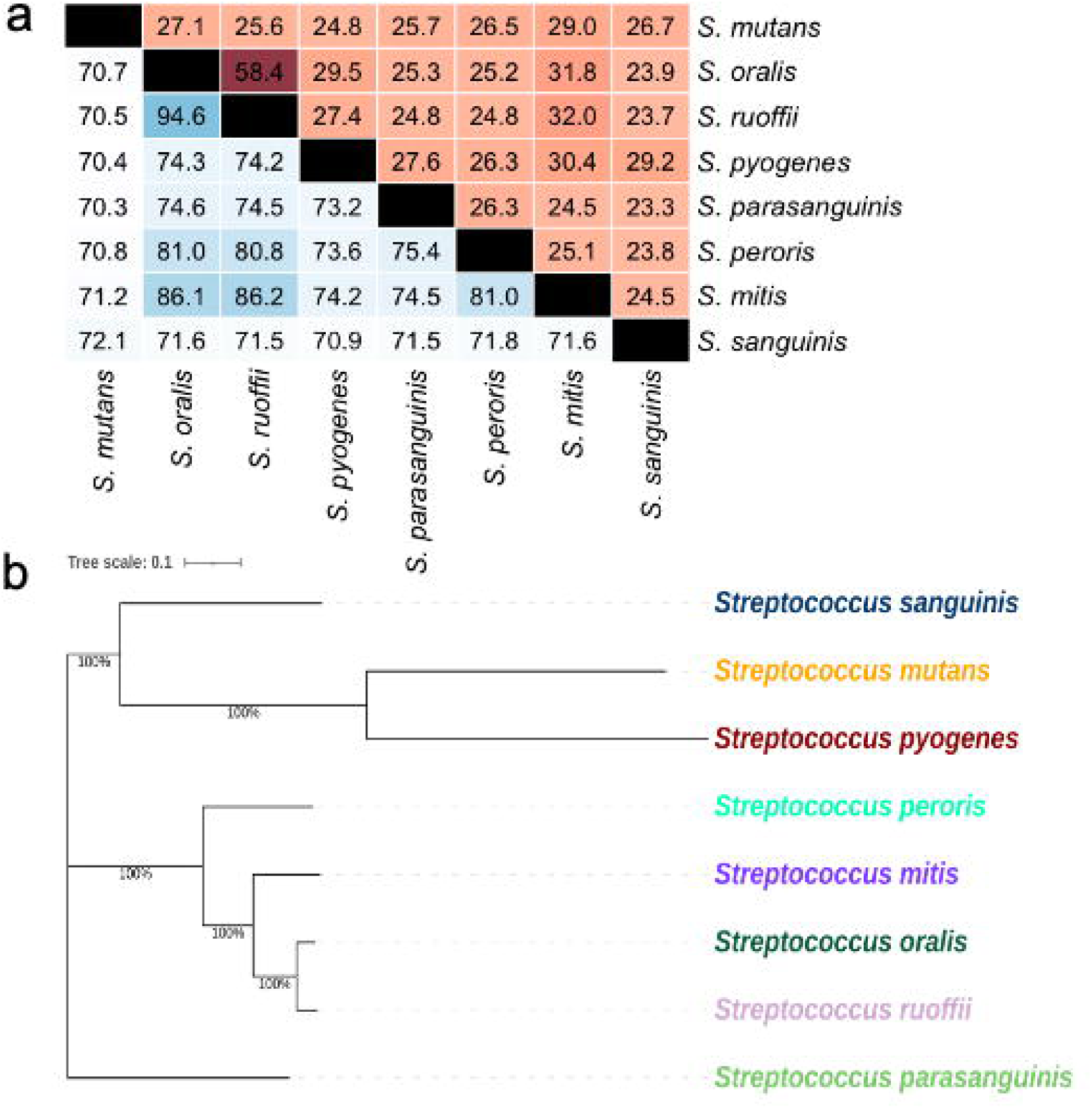
Phylogenetic analyses of *Streptococcus ruoffae* GP0011□. (a) Pairwise average nucleotide identity (ANI) values (top, blue-white gradient) and digital DNA–DNA hybridization (dDDH, d_4_) values (bottom, white-red gradient), between GP0011□ and the type strains of selected *Streptococcus* species. dDDH values calculated using the Genome-to-Genome Distance Calculator (GGDC, formula 2) on the Type Strain Genome Server (TYGS). ANI values calculated from the JSpeciesWS online service BLAST+ algorithm (ANIb). (b) Maximum likelihood phylogenomic tree based on 673 single-copy core genes, aligned with MAFFT and reconstructed in RAxML under the LG+F model of type strains of selected *Streptococcus* species. Strain designations and GenBank accession numbers can be found in Table 1. Branch values indicate bootstrap support (% from 100 replicates). Bar, 0.1 substitutions per site.

### Morphology and Growth of GP0011^T^

Measurements made using wet mount imaging showed ovoid-shaped cells with an average diameter of 1.3 µm arranged in chains of variable length (2-15 cells) and as individual cells (Fig. 2a-b). Gram-stained bacteria visualized by light microscopy confirmed the Gram-positive character of GP0011^T^. Imaging by scanning electron microscopy confirmed the morphology observed on the wet mounts (Fig. 2c). Growth on blood agar identified GP0011^T^ as α-hemolytic through assessment after 48 h incubation on trypticase soy broth (TSB) agar supplemented with 5% sheep blood (TBA) at 37 °C. Catalase activity was determined by applying 3% H_2_O_2_ to an overnight culture on a glass slide, confirming the absence of catalase for GP0011^T^. All the other *Streptococcus* species tested showed the consistent Gram nature, hemolysis, and catalase activity (Table 2).

**Fig. 2.**
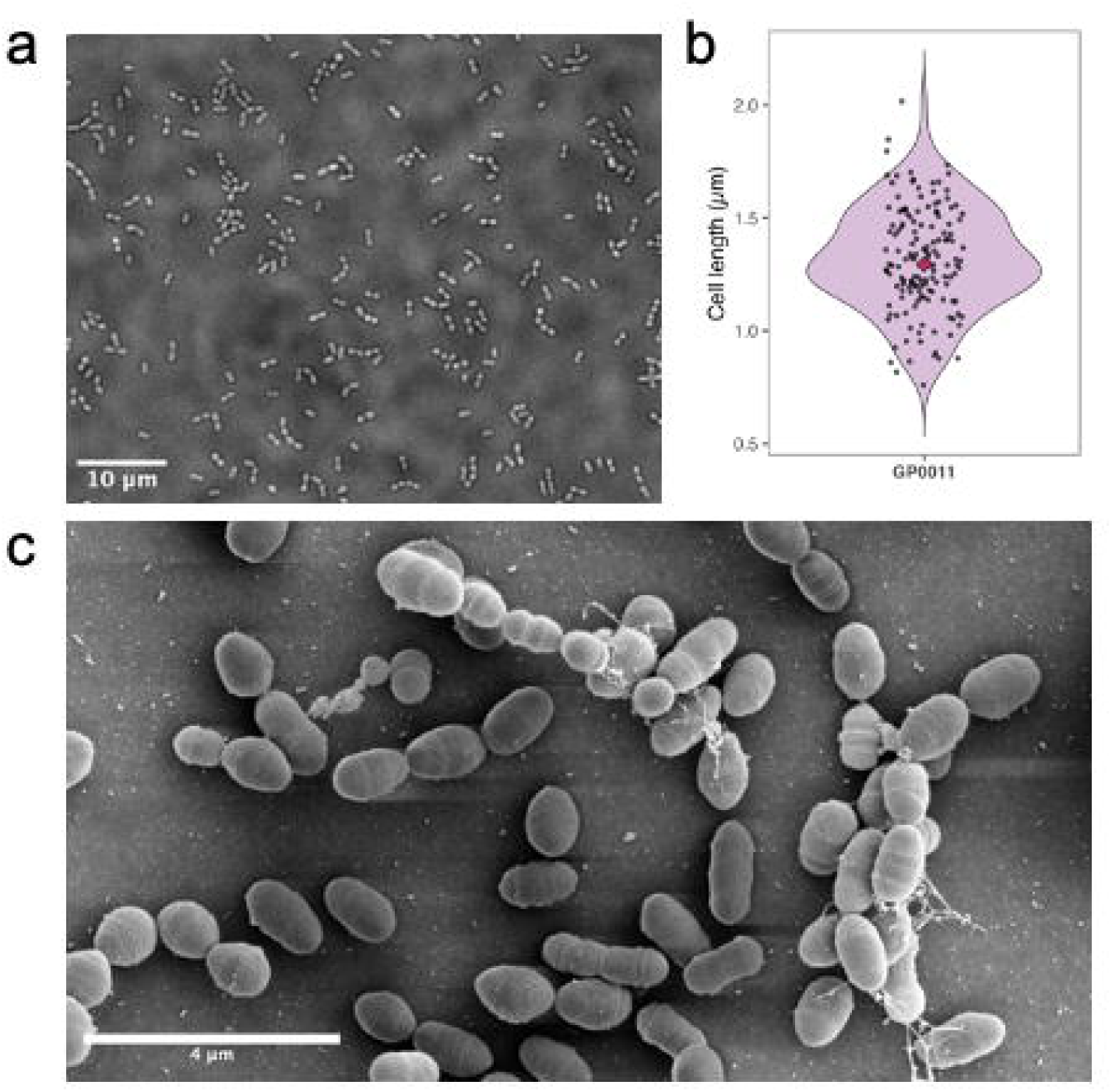
Morphological characterization of *Streptococcus ruoffae* GP0011□. (a) Light micrograph of GP0011□ cells prepared by wet mount with the Eclipse TI-E Inverted Research Microscope using 100x oil immersion objective. Bar, 10 µm. (b) Violin plot showing the distribution of cell diameter (µm) measured from wet mount images of GP0011□ ; the red diamond indicates the mean cell diameter (1.3 µm; n =150 between 3 biological replicates). (c) Scanning electron micrograph of GP0011□ cells with 21 pA Helios 5 at 25,000x magnification. Bar, 4 µm.

**Table 2.** Phenotypic characteristics of *Streptococcus ruoffae* GP0011□ and type strains of related *Streptococcus* species.

|  | ***S. ruoffae*** | ***S. oralis*** | ***S. mutans*** | ***S. sanguinis*** | ***S. parasanguinis*** | ***S. peroris*** | ***S. mitis*** | ***S. pyogenes*** |
|---|---|---|---|---|---|---|---|---|
| Strain | GP0011<sup>T</sup> | ATCC 35037<sup>T</sup> | ATCC 2517-5<sup>T</sup> | ATCC 10556<sup>T</sup> | ATCC 15912<sup>T</sup> | ATCC 700780<sup>T</sup> | OT25 | NCTC 12064 |
| Gram Staining | Gram-positive | Gram-positive | Gram-positive | Gram-positive | Gram-positive | Gram-positive | Gram-positive | Gram-positive |
| Morphology | Cocci in chains | Cocci in chains | Cocci in chains | Cocci in chains | Cocci in chains | Cocci in chains | Cocci in chains | Cocci in chains |
| Catalase | negative | negative | negative | negative | negative | negative | negative | negative |
| Hemolysis | α | α | α | α | α | α | α | ß |
| API20 Strep |  |  |  |  |  |  |  |  |
| VP |  |  | + |  |  |  |  |  |
| HIP |  |  |  |  |  |  |  |  |
| ESC: |  |  | + |  |  |  |  |  |
| PYRA |  |  |  |  |  |  |  | + |
| αGAL |  |  | + |  | + |  |  |  |
| ßGUR |  |  |  |  |  |  |  |  |
| ßGAL |  |  |  |  |  |  |  |  |
| PAL |  | + |  |  | + |  |  | + |
| LAP | + | + | + | + | + | + | + | + |
| ADH |  |  |  |  |  |  |  | + |
| RIB |  |  |  |  |  |  |  |  |
| ARA |  |  |  |  |  |  |  |  |
| MAN |  |  | + |  |  |  |  |  |
| SOR |  |  | + |  |  |  |  |  |
| LAC | + | + | + | + | + | + | + | + |
| TRE |  | + | + | + | + | + |  | + |
| INU |  |  | + |  |  |  |  |  |
| RAF |  |  | + | + | + |  |  |  |
| AMD |  | + |  |  |  |  |  |  |
| GLYG |  |  |  |  |  |  |  |  |

Legend: +, positive; –, negative. Hemolysis patterns on 5% sheep blood agar are indicated as α (alpha/partial hemolysis) or β (beta/complete hemolysis).

GP0011^T^ has a 1.34 h doubling time when grown aerobically on Todd-Hewitt Broth with 2.5% yeast supplemented (THY) medium with 5% CO_2_ at 37°C, and a 0.91 h doubling time anaerobically in comparison to 0.77 h and 0.52 h, respectively, for *S. oralis*. GP0011^T^ grew from 25-37°C aerobically (30°C optimum) and 25-43°C anaerobically (30°C optimum; Fig. 3a). GP0011^T^ grew from pH 6-10 aerobically (pH 8 optimum) and pH 5-10 anaerobically (pH 8 optimum). Growth occurred in NaCl from 0.0-2.0 g L^-1^ (optimum 0.5 g L^-1^) under both aerobic plus CO_2_ and anaerobic conditions (Fig. 3cd).

**Fig. 3.**
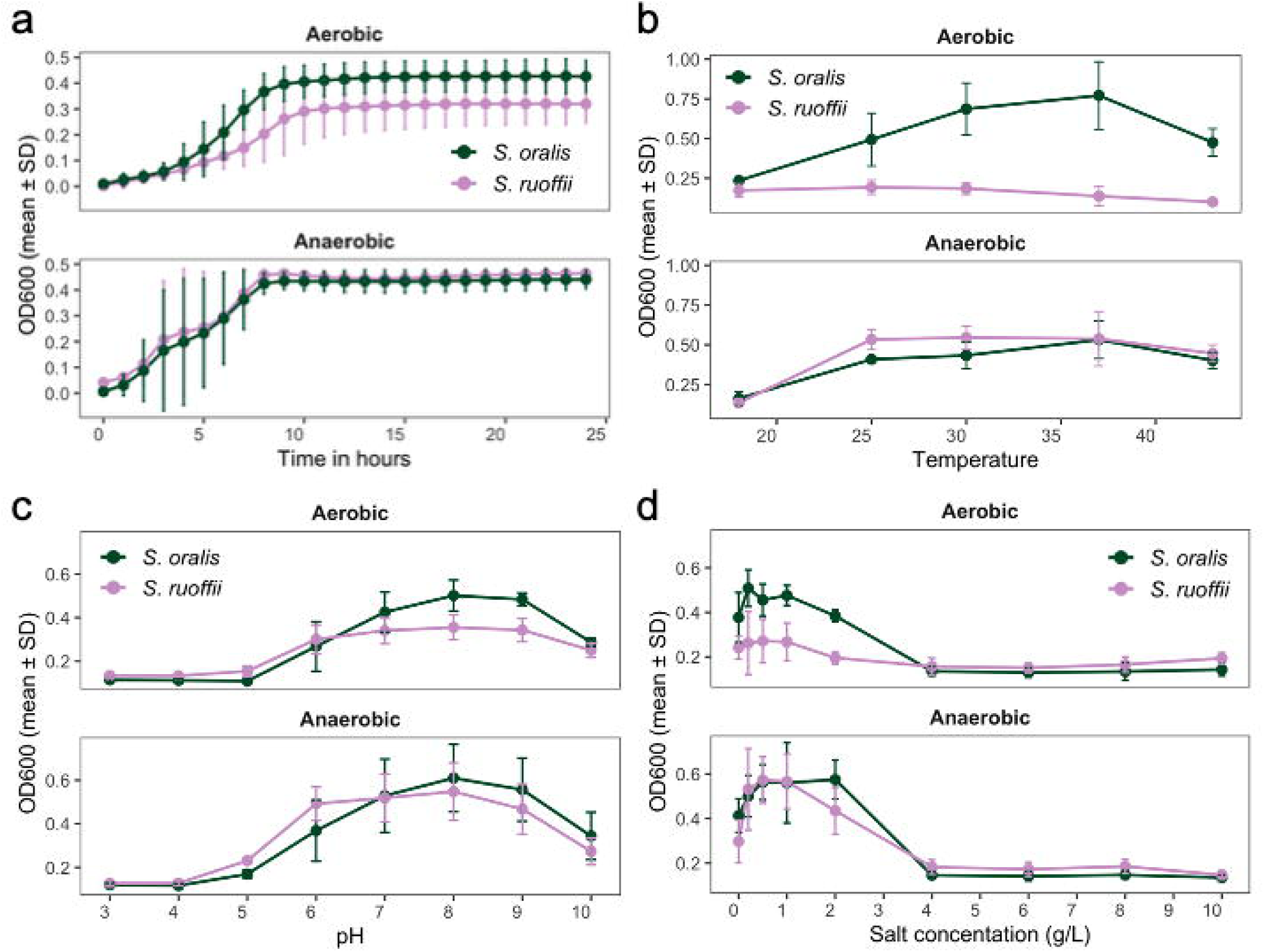
Physiological characterization of *Streptococcus ruoffae* GP0011□ (lilac) and *S. oralis* ATCC 35037□ (green). (a) Growth curves measured by OD_600_ over 24 h under aerobic with 5% CO_2_ (top) and anaerobic (bottom) conditions (anaerobic chamber with 2-3% H_2_ and 10% CO_2_) in THY broth; the fitted line represents the mean of three biological replicates, and shaded regions represent the confidence interval. (b) Terminal OD_600_ after 24 h exposure to temperatures from 18 to 43 °C under aerobic with 5% CO_2_ (top) and anaerobic (bottom) conditions in THY broth. (c) Terminal OD_600_ after 24 h at pH values from 3.0 to 10.0 under aerobic with 5% CO_2_ (top) and anaerobic (bottom) conditions in THY broth. (d) Terminal OD_600_ after 24 h across NaCl concentrations from 0 to 10 g l^-1^ under aerobic with 5% CO_2_ (top) and anaerobic (bottom) conditions in THY broth; for all panels points represent the mean of three biological replicates ±SD. The growth is defined as OD_600_>0.2.

### Distinct Biochemical Activities of GP0011^T^

Results indicated that GP0011^T^ and *S. oralis* could be distinguished by positive test results for alkaline phosphatase activity, trehalose fermentation, and starch fermentation in *S. oralis* only. Both species showed positive results in leucine aminopeptidase activity and lactose fermentation (Table 2). Several carbon sources allowed superior growth of *S. oralis* compared to that of GP0011^T^ under aerobic (Fig. 4a) and anaerobic (Fig. 4b) conditions. While none of the values under aerobic conditions were statistically significant, *S*. *oralis* showed OD_600_ of at least 0.2 higher when grown in the presence of sucrose, N-acetyl-β-D-mannosamine, N-acetyl-β-D-glucosamine, maltotriose, α-D-glucose, maltose, D-galactose, D-mannose, D-trehalose and α-D-lactose (Fig. 4c). Under anaerobic conditions, GP0011^T^ grew significantly better (P < 0.05) than *S*. *oralis* when the base medium was supplemented with D-mannitol and D-sorbitol. In contrast, S. *oralis* showed a positive trend in growth in comparison to GP0011^T^ when the base medium was supplemented with N-acetyl-β-D-mannosamine, D-trehalose, and lactulose (Fig. 4d). The superior growth of *S. oralis* with D-trehalose aligns with the positive API20 Strep test result observed for *S. oralis* compared to the negative test result for GP0011^T^ (Table 2).

**Fig. 4.**
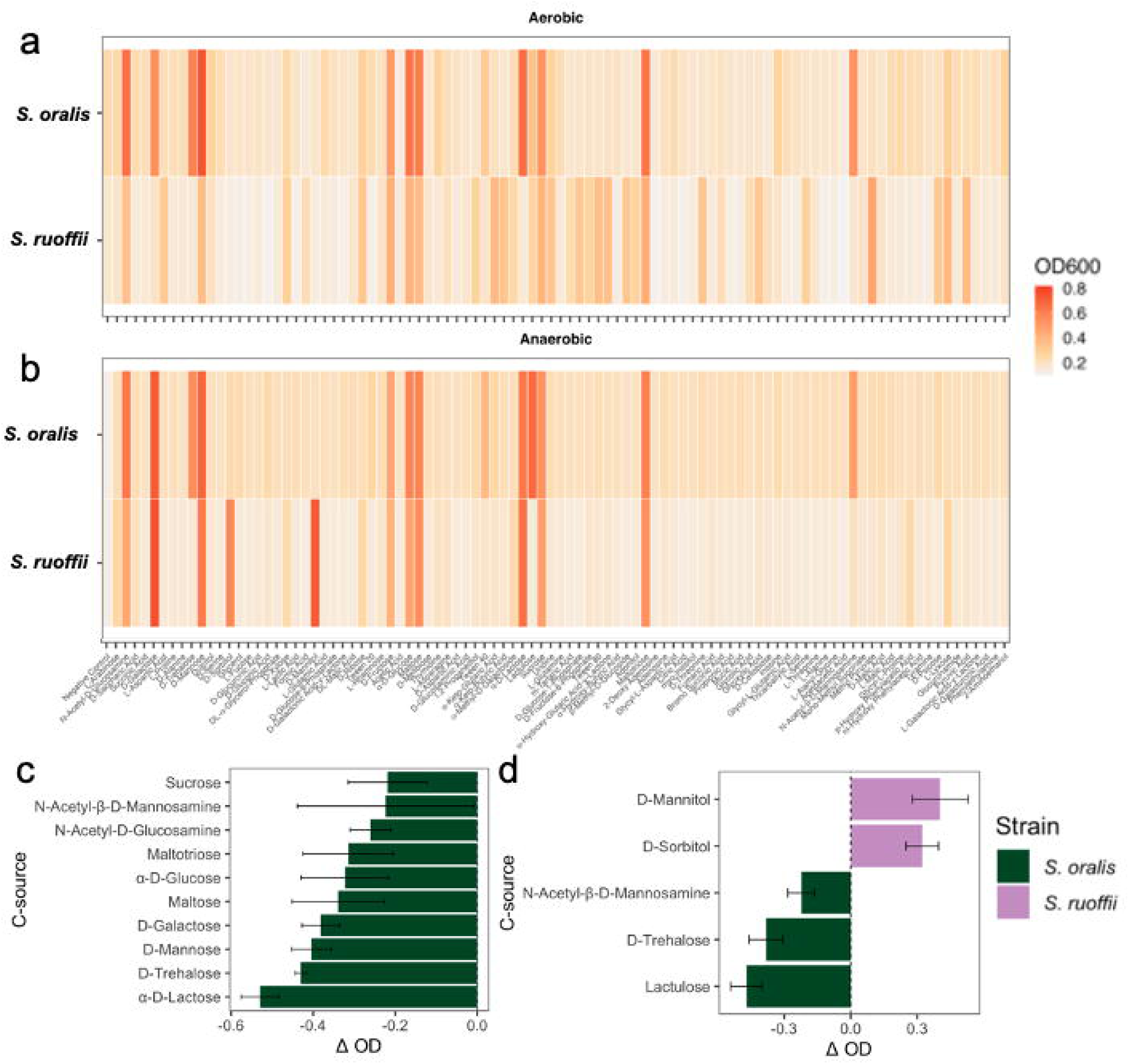
Biochemical characterization of *Streptococcus ruoffae* GP0011□ and *S. oralis* ATCC 35037□ via Biolog Phenotype MicroArray carbon utilization assays. (a) Heatmap of OD_600_ values for GP0011□ (*S. ruoffae*, bottom) and *S. oralis* (top) across various carbon sources under (a) aerobic and (b) anaerobic conditions after 24 h in minimal THY medium. The color scale indicates terminal OD_600_ after 24 h. Carbon sources with differential utilization between GP0011□ and *S. oralis* under (c) aerobic and (d) anaerobic conditions. Bars represent mean OD_600_ from three biological replicates; differences in aerobic conditions were identified with an OD_600_ difference > 0.2 between strains, and in anaerobic conditions with an OD_600_ difference> 0.2 between strains and with p < 0.05 (t-test, Benjamini–Hochberg correction). Positive values indicate higher utilization by *S. ruoffae* GP0011□ compared to *S. oralis* ATCC 35037□.

Using the BlastKOALA annotation tool, a pathway in the metabolism of trehalose was identified in the genome of *S. oralis*, but absent from the genome of GP0011^T^ [21]. Further supporting the differential metabolic capabilities of GP0011^T^ and *S. oralis*, five genes involved in mannitol and sorbitol transport and catabolism were identified as present in GP0011^T^ but absent from *S. oralis*. For mannitol catabolism, GP0011^T^ encodes a phosphotransferase (PTS) mannitol-specific transporter subunit IIBC, which imports mannitol as mannitol-1-phosphate via PTS-mediated phosphorylation during transport. This intermediate is then oxidized to fructose-6-phosphate by mannitol-1-phosphate 5-dehydrogenase (EC 1.1.1.17), enabling entry into glycolysis. For sorbitol, GP0011^T^ encodes all three subunits of the PTS glucitol/sorbitol transporter (IIA-IIC), which imports sorbitol as sorbitol-6-phosphate. Sorbitol-6-phosphate 2-dehydrogenase (EC 1.1.1.140) then oxidizes this intermediate to fructose-6-phosphate, similarly, feeding into glycolysis. These data are consistent with the ability of GP0011^T^ to grow on mannitol and sorbitol.

The major cellular fatty acids (>10.0% each) of GP0011^T^ were stearate (C18:0, 45.1%) and palmitate (C16:0, 31.1%); minor components included trans-9-elaidate (C18:1, 8.5%), arachidate (C20:0, 4.6%), palmitoleate (C16:1, 3.7%), heptadecanoate (C17:0, 3.7%), oleate (C18:1, 1.2%), and myristate (C14:0, 1.0%) (Fig. 5ab; S3, supplementary data). GP0011 was significantly enriched in stearate (C18:0) and palmitate (C16:0) relative to *S. oralis*, while *S. oralis* showed significantly higher composition of linoleate (C18:2), *cis*-11,14-eicosadienoate (C20:2), *cis*-10-heptadecenoate (C17:1), oleate (C18:1), *cis*-11-eicosenoate (C20:1), *trans*-9-elaidate (C18:1), tricosanoate (C23:0), palmitoleate (C16-1), and pentadecanoate (C15:0) compared to GP0011 (Fig. 5b; S3, supplementary data).

**Fig. 5.**
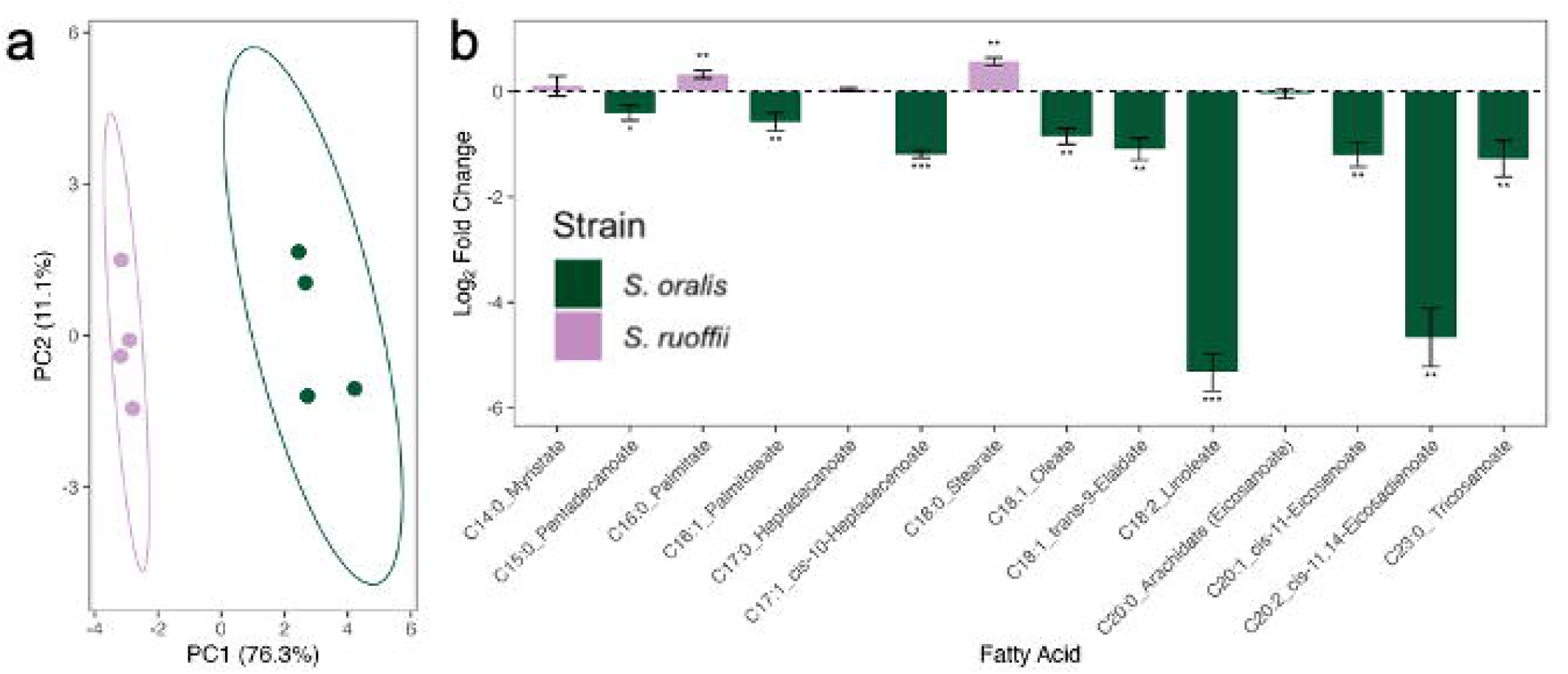
Cellular fatty acid profiles of *Streptococcus ruoffae* GP0011□ (lilac) and *S. oralis* ATCC 35037□ (green) determined by fatty acid methyl ester analysis (FAME) via GC-MS. (a) Principal component analysis (PCA) of normalized fatty acid profiles; each point represents one biological replicate (*n* = 4 per strain). Ellipses indicate 95% confidence intervals. (b) Log_2_ fold change of normalized fatty acid peak areas in *S. oralis* relative to GP0011□. Positive values indicate higher abundance in GP0011□; negative values indicate higher abundance in *S. oralis*. Bars represent mean ± SD (*n* = 4 biological replicates). Significance determined by Student’s t-test: \**p* < 0.05, \*\**p* < 0.01, \*\*\**p* < 0.001. The legend for both panels is shown in panel b.

### Antimicrobial Resistance Profiles of GP0011

*S. oralis* and GP0011^T^ displayed similar resistance profiles across the antibiotics tested, apart from GP0011^T^ having a minimum inhibitory concentration (MIC) for tetracycline of 1 µg mL^-1^ compared to the 4 µg mL^-1^ for *S. oralis* (Table 3). In comparison to *S. oralis*, GP0011^T^ was found to be more resistant to azithromycin, erythromycin, and tobramycin (Table 3). Using the ResFinder tool, from the Global Pathogens Analysis Platform, two antimicrobial resistance genes were identified in the GP0011^T^ genome: *msrD* (accession no. AF274302.1) and *mefA* (accession no. U83667.1), with 100% identity to the reference, which are not present in *S. oralis* [22]. The product of the *msrD* gene confers resistance to erythromycin, azithromycin, telithromycin, quinupristin, pristinamycin IA, and virginiamycin S, while carrying the *mefA* gene confers resistance to erythromycin and azithromycin. The resistance to erythromycin and azithromycin provided by these genes aligns with the antibiotic resistance assays described above.

**Table 3.** Minimum inhibitory concentrations (MICs; µg mL^-1^) for *Streptococcus ruoffae* **GP0011□ and *S. oralis* ATCC 35037□.**

| **Antibiotic** | ***S. ruoffae*** | ***S. oralis*** |
|---|---|---|
| Azithromycin (E) | 4 | 0.125 |
| Ciprofloxacin (E) | 1 | 1 |
| Clindamycin (E) | 0.032 | 0.016 |
| Erythromycin (E) | 3 | 0.016 |
| Ertapenem (E) | 0.003 | 0.006 |
| Levofloxacin (E) | 1.25 | 1 |
| Linezolid (E) | 0.5 | 0.25 |
| Piperacillin/Tazobactam (E) | 0.023 | 0 |
| Cefepime (E) | 0.064 | 0.047 |
| Tobramycin (E) | 128 | 32 |
| Trimethoprim/Sulfamethoxazole (E) | 0.19 | 0.125 |
| Amoxicillin/Clavulanate (E) | 0.016 | 0 |
| Carbenicillin (M) | 2 | 2 |
| Chloramphenicol (M) | 4 | 4 |
| Gentamicin (M) | >128 | 128 |
| Kanamycin (M) | >128 | >128 |
| Tetracycline (M) | 1 | 4 |

Legend: E-tests performed on TBA plates after 24 h (E); broth microdilution in THY after 24 h (M) using a 2-fold serial dilution series (128 to 0.0625 µg mL<sup>-1</sup>), following CLSI recommendations. Values represent the median of three biological replicates.

### Description of isolate with the proposed name *Streptococcus ruoffae* sp. nov

*Streptococcus ruoffae* (ru.off’i.i. N.L. gen. n. *ruoffae*, named in honor of Kathryn L. Ruoff for her contributions to the study of streptococci in the context of clinical microbiology). GP0011^T^ cells are Gram stain-positive, catalase-negative, facultatively aerobic ovoids, approximately 1-2 µm in diameter, occurring in short chains. After growth on TBA for 48 h at 37 °C with 5% CO_2_, colonies are yellow, opaque, circular, rounded, smooth, 0.5-1.0 mm in diameter, and α-hemolytic. The strain grows preferentially under anaerobic conditions. The temperature range for growth is 25-43 °C (optimum 30 °C) and the pH range for growth is 5.0-10.0 (optimum pH 8.0). The strain grows in broth medium with 0.0-2.0 g L^-1^ NaCl (optimum 0.5 g L^-1^ NaCl). Using API 20 STREP, positive reactions are observed for lactose fermentation and leucine aminopeptidase activity. Mannitol and sorbitol were identified as the preferred carbon sources for *S. ruoffae*, and analysis of the genome identified genes coding for components of the catabolic pathways. The major cellular fatty acids are stearate and palmitate. Antimicrobial resistance genes for erythromycin and azithromycin were detected in the genome, and the resistance was confirmed phenotypically.

The type strain was isolated from a human endoscopy sample of an adult male CF heterozygote in Lebanon, New Hampshire, United States. The DNA G+C content of the type strain is 40.93%. The GenBank accession numbers for the genome and 16S rRNA sequence are GCA_030546305.1 and PZ058466, respectively. The type strain is GP0011^T^ (ATCC TSD-627 and NCTC 15312).

## DISCUSSION

The characterization of GP0011^T^ supports its designation as a novel species within the *Streptococcus* mitis group [3]. 16S rRNA gene sequence analysis alone placed GP0011^T^ close to established mitis group members, but genome-based analyses gave a clearer picture. The highest dDDH value for GP0011^T^ was 58.3% with *S. oralis*, below the accepted species delineation threshold [17]. Its ANI with *S. oralis*, 94.84%, is consistent with that conclusion, falling below the 95-96% range generally applied for species delineation [23]. The core-genome phylogeny, built from 673 single-copy genes, placed GP0011^T^ as a distinct, well-supported lineage within the mitis group, providing independent support for the dDDH and ANI results. Members of the mitis group are often indistinguishable by 16S rRNA gene sequence alone [24, 25], and GP0011^T^ follows that pattern, sharing over 99% 16S identity with both *S. mitis* and *S. oralis* while remaining clearly distinct from each in genome-wide measures. These results support combining genome-level analyses with 16S rRNA sequencing for species delineation in the mitis group.

Phenotypic data provide additional support for this distinction. GP0011^T^ and *S. oralis* differed in several API20 Strep reactions and in the carbon sources each preferred. *S. oralis* grew better on D-trehalose, consistent with its positive API20 trehalose reaction, whereas GP0011^T^ was negative for this reaction and lacks the trehalose metabolism pathway identified in *S. oralis.* GP0011^T^, in turn, grew significantly better than *S. oralis* on D-mannitol and D-sorbitol under anaerobic conditions, and its genome carries the transport and catabolism genes for both sugars that *S. oralis* lacks. However, the D-sorbitol phenotype was inconsistent with the API20 result. GP0011^T^ grew on D-sorbitol anaerobically despite testing negative on the API20 sorbitol reaction. Both pathways proceed through phosphotransferase system transporters, which phosphorylate their substrate during import, followed by dehydrogenase-mediated conversion to fructose-6-phosphate that enters glycolysis; these systems have been well characterised in *S. mutans* [26, 27]. Commercial biochemical panels have been shown to perform poorly for species-level identification within the viridans group, with difficulty distinguishing *S. mitis* and *S. oralis* [28]. Additionally, the API20 Strep format is not performed under strict anaerobic conditions. Either could account for this observed disagreement. Overall, the agreement between genomic and phenotypic data suggests that these metabolic differences represent genuine distinctions in the metabolic capabilities of the two species rather than differences detectable only at the sequence level.

The preferential growth of GP0011^T^ anaerobically adds further insight into its physiology. GP0011^T^ had a shorter doubling time anaerobically than aerobically and grew across a broad range of temperatures and pH, with optima at 30 °C and pH 8. Shifts toward anaerobic and microaerophilic taxa have been described in the esophagus under inflammatory and metaplastic conditions [6, 7]. Its more robust growth under anaerobic conditions, along with its differential carbon source utilization under those conditions, suggests that *S. ruoffae* may be well suited to environments in which oxygen availability is limited. However, this study does not establish the specific environmental conditions supporting *S. ruoffae* in vivo; its physiological properties should not be interpreted as direct evidence of adaptation to the esophagus.

The cellular fatty acid profile provides an additional chemotaxonomic distinction between *S. ruoffae* and *S. oralis*. GP0011^T^ was enriched in stearate and palmitate, whereas *S. oralis* contained higher proportions of several other fatty acids, a pattern consistent with the major fatty acids reported for other recently described mitis-group-adjacent species, including *S. jiangnanensis* [29], one of the comparator strains used in this phylogenetic analysis. Together with the physiological and genomic differences observed between the strains, these chemotaxonomic differences provide independent phenotypic evidence supporting the separation of GP0011^T^ from its closest relative.

The antimicrobial resistance profile of *S. ruoffae* is also notable. It carries the acquired resistance genes *msrD* and *mefA*, absent from *S. oralis*, and is phenotypically more resistant to erythromycin and azithromycin. In streptococci, *mefA* and *msrD* occur as an adjacent gene pair on mobile elements such as Tn*1207.1* and Φ1207.3 [30], with *msrD* encoding the ABC-transporter component required for resistance and *mefA* contributing a smaller effect [31]. Their presence therefore represents potentially transferable resistance rather than a chromosomal trait restricted to this isolate.

The correspondence between the presence of the *msrD* and *mefA* genes and observed resistance phenotype is consistent with their contribution to the antimicrobial resistance profile of GP0011^T^. The mitis group has long been recognized as a reservoir of resistance determinants acquired by *S. pneumoniae* through natural transformation and homologous recombination [32].

Mobile macrolide resistance genes in a commensal from this group are therefore of interest beyond an individual isolate. Azithromycin is used as long-term maintenance therapy in cystic fibrosis, therefore macrolide resistance determinants in organisms recovered from individuals with CF-associated genotypes are of potential clinical interest [12, 13, 33]. However, the isolation of a single resistant strain from a CFTR heterozygote does not establish an association between *S. ruoffae*, CFTR heterozygosity, or antimicrobial exposure.

The recovery of *S. ruoffae* from the esophagus expands the known species-level diversity of the esophageal microbiome. *Streptococcus* is a predominant genus in the healthy esophageal microbiome [5], yet most surveys of this site have characterised communities at the genus level [5, 6, 7]. Culture-based approaches continue to recover previously undescribed taxa from human body sites [34]. The identification and genomic characterization of GP0011^T^ therefore provides a reference genome and type strain that can facilitate recognition of this taxon in future studies. In particular, the combination of genomic and phenotypic characteristics described here may help distinguish *S. ruoffae* from closely related *Streptococcus* species in culture-based and sequence-based investigations.

Several limitations should be considered when interpreting these findings. The description of *S. ruoffae* is based on a single isolate obtained from one individual; therefore, the prevalence and distribution of this species cannot be determined from this study. Similarly, although the isolate was recovered from a CFTR heterozygote and carriers have been reported to show subclinical mucosal phenotypes and elevated infection risk [12, 13, 14], these data cannot determine whether *S. ruoffae* is associated with CF, CFTR heterozygosity, or any particular clinical condition. Additional isolates from individuals with different clinical backgrounds and from other anatomical sites would be necessary to determine the distribution and diversity of the species. Future work examining additional *S. ruoffae* isolates could help determine whether the characteristics observed in GP0011^T^ are representative of the species as a whole.

Overall, the genomic, phylogenetic, physiological, metabolic, chemotaxonomic, and antimicrobial susceptibility data distinguish GP0011^T^ from previously described *Streptococcus* species and support the proposal of *Streptococcus ruoffae* sp. nov. The identification of this species from the esophagus highlights the continued potential for species-level discovery within relatively understudied human microbial niches [4, 34] and provides a genomic and phenotypic reference for future investigations of *Streptococcus* diversity.

## MATERIALS AND METHODS

Whole-genome sequencing of GP0011^T^ has been published [15] and can be accessed using accession GCA_030546305.1. The digital DNA–DNA hybridization (dDDH) values were calculated using Genome-to-Genome Distance (GGDC, formula 2) on the type strain genome server (TYGS) [16]. To further establish the phylogenetic relationship of strain GP0011^T^ to other members of the genus, seven other species were chosen for comparative analysis in this study (Table 1). These strains were analysed for dDDH and average nucleotide identity (ANI) analysis. The phylogenetic tree based on the 16S rRNA gene was generated for the selected species by RAxML using the GTR-GAMMA model. Because species within the mitis group often cannot be reliably differentiated on the basis of 16S rRNA gene identity alone [24, 25], additional phylogenetic analyses were performed with housekeeping genes *rpoB* and *gyrB* [19].

Samples for this study were collected following institutional protocols approved by Dartmouth Health (IRB #07876). One former smoker between 60-65 years of age, a heterozygote for the CF allele provided an esophageal sample from an endoscopy procedure [34]. To isolate bacteria, the samples were cultured on TBA, *Bacteroides* bile esculin agar (BBE) and MacConkey agar (McC) media. Cultures were incubated for 48 h at 37 °C under both aerobic and anaerobic conditions, with some of the plates supplemented with 100 µg mL^-1^ gentamicin as a selective agent. Individual colonies were then purified through three rounds of streaking. From this sample, we isolated GP0011^T^. Unless otherwise stated, GP0011^T^ was grown in THY at 37 °C for phenotypic characterization and anaerobic growth conditions were attained via the GasPak EZ Anaerobe Container Systems (Becton Dickinson, USA).

Cell morphology, size, and Gram stain reaction were examined with light microscopy or scanning electron microscopy (SEM), as indicated. Light microscopy used the Eclipse TI-E Inverted Research Microscope (Nikon, Japan) to observe cells using a standard wet mount with 1% agarose pad. For scanning electron microscopy (SEM), cells were grown overnight in liquid culture on glass coverslips, were negatively stained, and imaged using the Helios 5 DualBeam (Thermo Fisher Scientific, United States) SEM. The standard Gram stain procedure was performed with the BD Gram Staining Kit by heat-fixing cells and staining.

Growth curves were obtained using the Stratus Miniaturized Plate Reader (Cerillo, USA) under aerobic condition with 5% CO_2_ and with the use of an anaerobic chamber (Coy, USA) with 3% H_2_ and 10% CO_2_ [35]. Optical density measurements at 600 nm were obtained at 3-minute intervals for 24 h, comparing GP0011^T^ and *S. oralis* growth patterns. Growth statistics were determined with the R package growthcurver [36].

Growth at a range of temperatures (18°C, 25°C, 30°C, 37°C, 43°C) was assessed in THY aerobically plus 5% CO_2_ and anaerobically after 24 h. Similarly, pH values ranging from 3.0-10.0 (at intervals of 1.0 pH units) and various NaCl concentrations (0.0, 0.2, 0.5, 1.0, 2.0, 4.0, 6.0, 8.0, 10.0 g L^-1^) were also assessed in THY aerobically plus 5% CO_2_ and anaerobically after 24 h.

Biochemical characterization was performed using the API 20 STREP system (bioMérieux, France). Tests were performed in parallel with the seven other species described here. API20 STREP profiles for each of the tested species matched with the manual identification table provided by the manufacturer. To determine the carbon sources utilized by GP0011^T^, we utilized a base medium composed of THY broth without glucose. This medium was used to test growth of GP0011^T^ and *S. oralis* in various carbon sources using Biolog Phenotype MicroArray Carbon Utilization Assay Plates (Biolog, USA) under aerobic plus 5% CO_2_ or anaerobic conditions (Table S1-2, supplementary data). GP0011^T^ cells grown in brain-heart infusion (BHI) broth at 37°C for 24 h under 5% CO_2_ were subjected to cellular fatty acid analysis by Fatty Acid Methyl Ester Analysis (FAME) through gas chromatography-mass spectrometry (GC-MS) using an Agilent 8890 GC-MS at the UMass Chan Metabolomics Core.

Phenotypic antimicrobial susceptibility was determined using E-TEST (bioMérieux, France) strips and using the broth microdilution method according to Clinical and Laboratory Standards Institute (CLSI M100 standards, 2020) recommendations. The broth microdilution method involved a 2:1 serial dilution of the indicated antibiotic in liquid culture from 128 µg mL^-1^ to 0.0625 µg mL^-1^ for a series of antibiotics; the bacteria were grown for 24 h in THY. We also assessed antimicrobial activity using the E-TEST strips with bacteria grown on TBA medium plates for 24 h [37].

## Acknowledgments

The authors thank Kathryn Ruoff for her decades of service working with *Streptococcus* species and her years working in the O’Toole lab. We would like to thank Radu Stan (Dartmouth Electron Microscopy Facility) and Joseph Schwartzman for their respective assistance with imaging and analysis. We would also like to thank Nicholas Jacobs for providing the *S. mitis* and *S. pyogenes* strains tested. We thank the UMass Chan Medical School Metabolomics Core Facility (RRID:SCR_027036) for GC-MS analysis.

## Data Availability

The strain is available from ATCC TSD-627 and NCTC 15312. The code and raw data used for analysis are available upon request. The whole-genome sequence for strain GP0011^T^ (*Streptococcus ruoffae sp. nov.*) was deposited in GenBank with the accession numbers GCA_030546305.1 for the genome and PZ058466 for the 16S rRNA gene sequence.

## References

1. Jorgensen JH, Pfaller MA, Carroll KC, Funke G, Landry ML, Richter SS, Warnock DW (ed). Manual of Clinical Microbiology. Washington, DC, USA: ASM Press; 2015.

2. Hardie JM, Whiley RA. Classification and overview of the genera *Streptococcus* and *Enterococcus*. J Appl Microbiol 1997;83:1S–11S.

3. Patel S, Gupta RS. Robust demarcation of fourteen different species groups within the genus *Streptococcus* based on genome-based phylogenies and molecular signatures. Infect Genet Evol J Mol Epidemiol Evol Genet Infect Dis 2018;66:130–151.

4. Freese HM, Meier-Kolthoff JP, Sardà Carbasse J, Afolayan AO, Göker M. TYGS and LPSN in 2025: a global core biodata resource for genome-based classification and nomenclature of prokaryotes within DSMZ Digital Diversity. Nucleic Acids Res 2026;54:D884–D891.

5. Bonazzi E, Lorenzon G, Maniero D, De Barba C, Bertin L, Barberio B, Salvador R, Valmasoni M, Zingone F, Ghisa M, Savarino EV. The esophageal microbiota in esophageal health and disease. Gastroenterol Insights 2024;15:998–1013.

6. Yang L, Lu X, Nossa CW, Francois F, Peek RM, Pei Z. Inflammation and intestinal metaplasia of the distal esophagus are associated with alterations in the microbiome. Gastroenterology 2009;137:588–597.

7. Baumeister T, Quante M, Bornschein J. Gut microbiota and oesophageal disease – an update. Microb Health Dis 2020;2:e207.

8. Cutting GR. Cystic fibrosis genetics: from molecular understanding to clinical application. Nat Rev Genet 2015;16:45–56.

9. Caley LR, White H, de Goffau MC, Floto RA, Parkhill J, Marsland B, Peckham DG. Cystic fibrosis-related gut dysbiosis: a systematic review. Dig Dis Sci 2023;68:1797–1814.

10. Burke DG, Fouhy F, Harrison MJ, Rea MC, Cotter PD, O’Sullivan O, Stanton C, Hill C, Shanahan F, Plant BJ, Ross RP. The altered gut microbiota in adults with cystic fibrosis. BMC Microbiol 2017;17:58.

11. Antosca KM, Chernikova DA, Price CE, Ruoff KL, Li K, Guill MF, Sontag NR, Morrison HG, Hao S, Drumm ML, MacKenzie TA, Dorman DB, Feenan LM, Williams MA, Dessaint J, Yuan IH, Aldrich BJ, Moulton LA, Ting L, Martinez-del Campo A, Stewart EJ, Karagas MR, O’Toole GA, Madan JC. Altered stool microbiota of infants with cystic fibrosis shows a reduction in genera associated with immune programming from birth. J Bacteriol 2019;201:e00274–19.

12. Polgreen PM, Brown GD, Hornick DB, Ahmad F, London B, Stoltz DA, Comellas AP. CFTR heterozygotes are at increased risk of respiratory infections: a population-based study. Open Forum Infect Dis 2018;5:ofy219.

13. Polgreen PM, Comellas AP. Clinical phenotypes of cystic fibrosis carriers. Annu Rev Med 2022;73:563–574.

14. Kalyta K, Stelmaszczyk W, Szczęśniak D, Kotuła L, Dobosz P, Mroczek M. The spectrum of the heterozygous effect in biallelic Mendelian diseases—the symptomatic heterozygote issue. Genes 2023;14:1562.

15. Surve SV, Valls RA, O’Toole GA. Draft genome sequences of 109 gut bacteria isolated from colonoscopy samples of individuals with and without cystic fibrosis. Microbiol Resour Announc 2026;0:e01114–25.

16. Meier-Kolthoff JP, Göker M. TYGS is an automated high-throughput platform for state-of-the-art genome-based taxonomy. Nat Commun 2019;10:2182.

17. Auch AF, Von Jan M, Klenk H-P, Göker M. Digital DNA-DNA hybridization for microbial species delineation by means of genome-to-genome sequence comparison. Stand Genomic Sci 2010;2:117–134.

18. Zhang Z, Schwartz S, Wagner L, Miller W. A greedy algorithm for aligning DNA sequences. J Comput Biol 2000;7:203–214.

19. Stamatakis A. RAxML version 8: a tool for phylogenetic analysis and post-analysis of large phylogenies. Bioinformatics 2014;30:1312–1313.

20. Katoh K, Misawa K, Kuma K, Miyata T. MAFFT: a novel method for rapid multiple sequence alignment based on fast Fourier transform. Nucleic Acids Res 2002;30:3059–3066.

21. Kanehisa M, Sato Y, Morishima K. BlastKOALA and GhostKOALA: KEGG tools for functional characterization of genome and metagenome sequences. J Mol Biol 2016;428:726– 731.

22. Zankari E, Hasman H, Cosentino S, Vestergaard M, Rasmussen S, Lund O, Aarestrup FM, Larsen MV. Identification of acquired antimicrobial resistance genes. J Antimicrob Chemother 2012;67:2640–2644.

23. Richter M, Rosselló-Móra R. Shifting the genomic gold standard for the prokaryotic species definition. Proc Natl Acad Sci U S A 2009;106:19126–19131.

24. Janda JM, Abbott SL. 16S rRNA gene sequencing for bacterial identification in the diagnostic laboratory: pluses, perils, and pitfalls. J Clin Microbiol 2007;45:2761–2764.

25. Glazunova OO, Raoult D, Roux V. Partial sequence comparison of the rpoB, sodA, groEL and gyrB genes within the genus *Streptococcus*. Int J Syst Evol Microbiol 2009;59:2317–2322.

26. Honeyman AL, Curtiss R 3rd. Isolation, characterization, and nucleotide sequence of the *Streptococcus mutans* mannitol-phosphate dehydrogenase gene and the mannitol-specific factor III gene of the phosphoenolpyruvate phosphotransferase system. Infect Immun 1992;60:3369– 3375.

27. Boyd DA, Thevenot T, Gumbmann M, Honeyman AL, Hamilton IR. Identification of the operon for the sorbitol (glucitol) phosphoenolpyruvate:sugar phosphotransferase system in *Streptococcus mutans*. Infect Immun 2000;68:925–930.

28. Kikuchi K, Enari T, Totsuka K, Shimizu K. Comparison of phenotypic characteristics, DNA-DNA hybridization results, and results with a commercial rapid biochemical and enzymatic reaction system for identification of viridans group streptococci. J Clin Microbiol 1995;33:1215– 1222.

29. Zhou Y, Zhao J, Ross P, Stanton C, Chen W, Yang B. *Streptococcus wuxiensis* sp. nov., *Streptococcus jiangnanensis* sp. nov. and *Streptococcus fermentans* sp. nov.: three novel species of genus *Streptococcus* isolated from human breast milk. Int J Syst Evol Microbiol 2025;75:006674. doi:10.1099/ijsem.0.006674

30. Santagati M, Iannelli F, Oggioni MR, Stefani S, Pozzi G. Characterization of a genetic element carrying the macrolide efflux gene *mef*(A) in *Streptococcus pneumoniae*. Antimicrob Agents Chemother 2000;44:2585–2587.

31. Iannelli F, Santoro F, Santagati M, Docquier J-D, Lazzeri E, Pastore G, Cassone M, Oggioni MR, Rossolini GM, Stefani S, Pozzi G. Type M resistance to macrolides is due to a two-gene efflux transport system of the ATP-binding cassette (ABC) superfamily. Front Microbiol 2018;9:1670.

32. Salvadori G, Junges R, Morrison DA, Petersen FC. Competence in *Streptococcus pneumoniae* and close commensal relatives: mechanisms and implications. Front Cell Infect Microbiol 2019;9:94.

33. Saiman L, Marshall BC, Mayer-Hamblett N, Burns JL, Quittner AL, Cibene DA, Coquillette S, Fieberg AY, Accurso FJ, Campbell PW 3rd. Azithromycin in patients with cystic fibrosis chronically infected with *Pseudomonas aeruginosa*: a randomized controlled trial. JAMA 2003;290:1749–1756.

34. Surve SV, Valls RA, Barrack KE, Gwilt LL, Gardner TB, O’Toole GA. The regional landscape of the human colon culturome in health and cystic fibrosis. Microbiol Spectr 2026;14:e03126–25.

35. Rogers AT, Bullard KR, Dod AC, Wang Y. Bacterial growth curve measurements with a multimode microplate reader. Bio-Protoc 2022;12:e4410.

36. Sprouffske K, Wagner A. Growthcurver: an R package for obtaining interpretable metrics from microbial growth curves. BMC Bioinformatics 2016;17:172.

37. Hazbón MH, del Socorro Orozco M, Labrada LA, Tovar R, Weigle KA, Wanger A. Evaluation of Etest for susceptibility testing of multidrug-resistant isolates of *Mycobacterium tuberculosis*. J Clin Microbiol 2000;38:4599–4603.

