## Supplemental Figures for "Identification and characterization of a *Streptococcus* strain isolated from the esophagus of an adult male heterozygous for the CF allele"

### Supplementary Figures

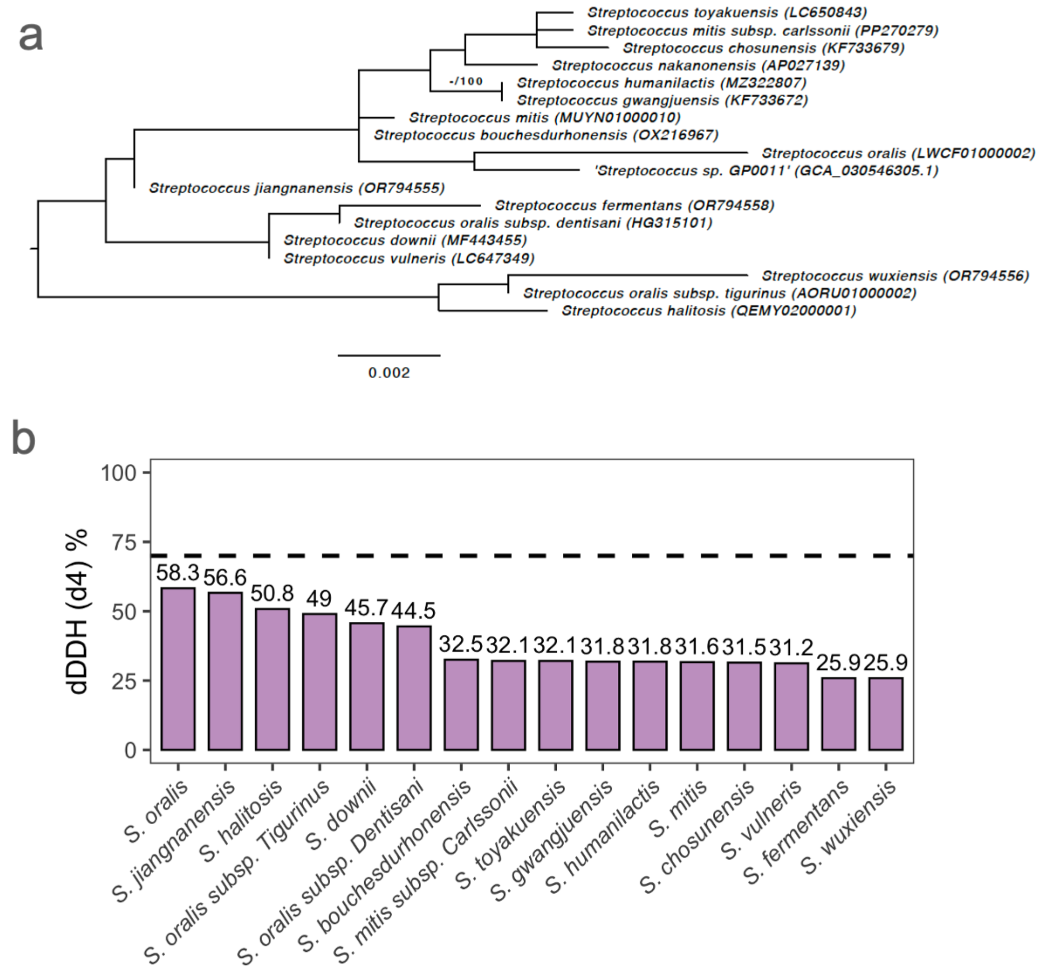


**Fig. S1.** Relationships between *Streptococcus ruoffae* GP0011$^{T}$ and the closest related streptococcal species. (a) Whole-genome phylogeny of *S. ruoffae* GP0011$^{T}$ and related *Streptococcus* type strains. Tree generated using the Type Strain Genome Server (TYGS) with the Genome BLAST Distance Phylogeny (GBDP) method. GenBank accession numbers given in parentheses. Bar, 0.002 substitutions per site. (b) Digital DNA–DNA hybridization (dDDH) values between *Streptococcus* GP0011$^{T}$ and its closest relatives, calculated using the Genome-to-Genome Distance Calculator (GGDC 3.0, formula $d_4$). The dashed line indicates the 70% species-delineation threshold.

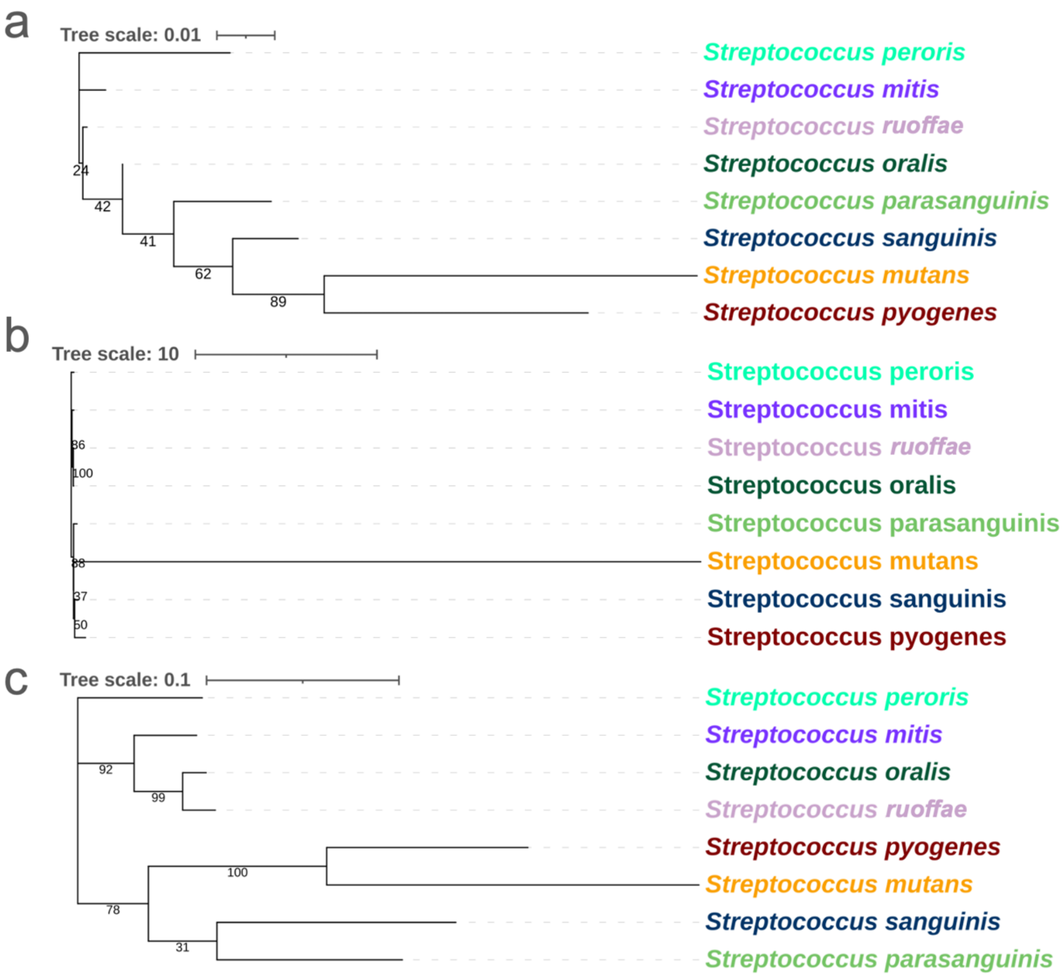


**Figure S2.** Maximum likelihood trees based on (a) 16S rRNA gene sequences, (b) *gyrB* (1946 bp), and (c) *rpoB* (3611 bp). Branch support values are resampling estimated log-likelihoods (RELL, %) and are shown at nodes. Bars indicate the respective substitutions per site as shown.

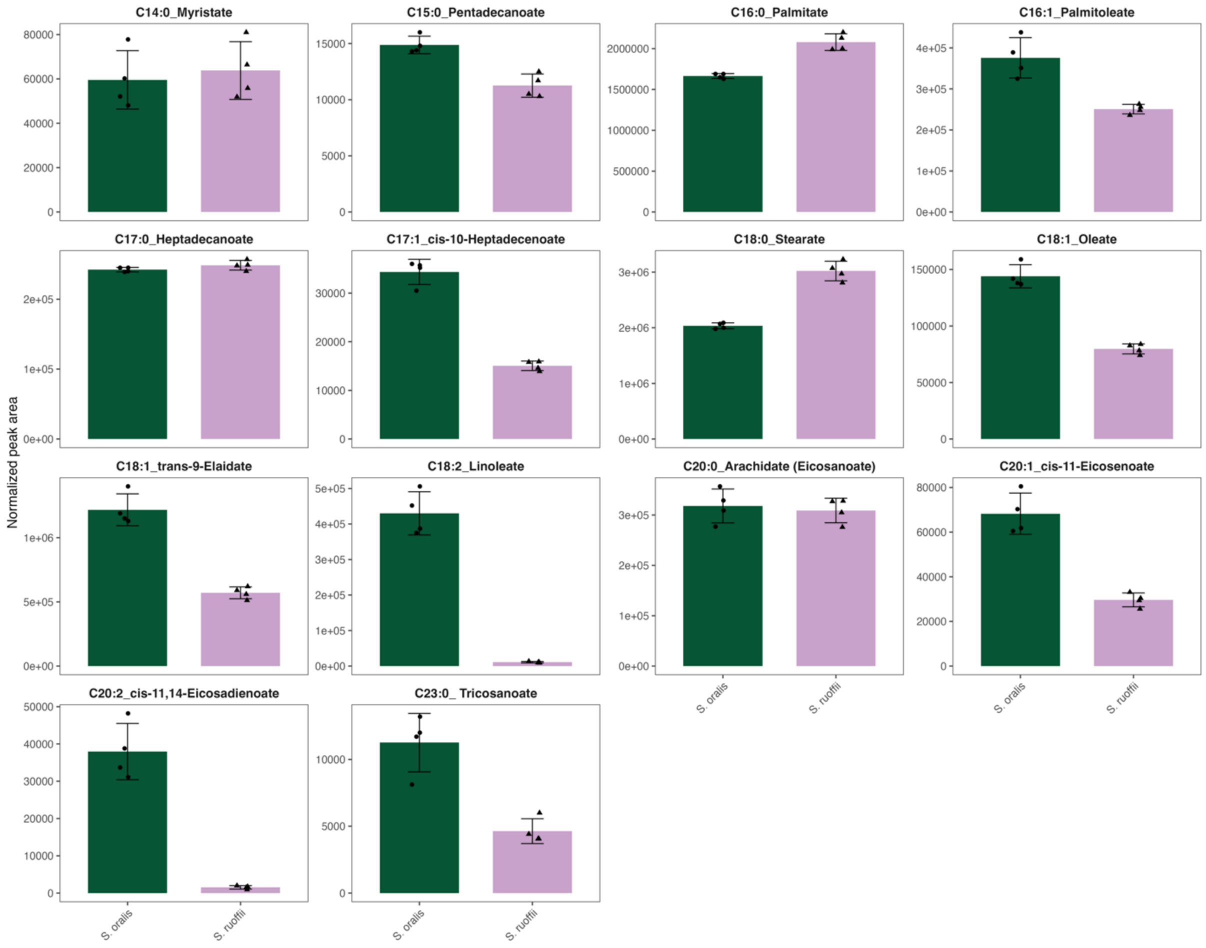


**Fig. S3.** Normalized fatty acid peak areas for *Streptococcus ruoffae* GP0011$^{T}$ (lilac) and *S. oralis* ATCC 35037$^{T}$ (green) across detected fatty acid species. Areas were normalized to a $^{13}C_1$-palmitate internal standard. Bars represent mean ± SD (*n* = 4 biological replicates).
